# Elevational gradients do not affect thermal tolerance at local scale in populations of livebearing fishes of the genus *Limia* (Teleostei, Poeciliidae)

**DOI:** 10.1101/2020.12.26.424431

**Authors:** Rodet Rodriguez Silva, Ingo Schlupp

## Abstract

One of the main assumptions of Janzen’s (1976) mountain passes hypothesis is that due the low overlap in temperature regimes between low and high elevations in the tropics, organisms living in high-altitude evolve narrow tolerance for colder temperatures while low-altitude species develop narrow tolerance for warmer temperatures. Some studies have questioned the generality of the assumptions and predictions of this hypothesis suggesting that other factors different to temperature gradients between low and high elevations may explain altitudinal distribution of species in the tropics. We assessed variation in tolerance to extreme temperatures (measured as critical thermal minimum (CTmin) and maximum (CTmax)) and also compared thermal breadth for populations of eight species of livebearing fishes of the genus *Limia* occurring in three Caribbean islands and that occupy different altitudinal distribution. Our results showed that species analyzed had significant differences in thermal limits and ranges. Generally, species distributed in high and low elevations did not differ in thermal limits and showed a wider range of thermal tolerance. However, species living in mid-elevations had narrower range of temperature tolerance. We found no significant effect of phylogeny on CTmin, CTmax and thermal ranges among species. This study did not provide evidence supporting Janzen’s hypothesis at a local scale since thermal tolerance and altitudinal distribution of *Limia* species were not related to temperature gradients expected in nature. Phylogeny also did not explain the patterns we observed. We suggest that biotic factors such as species interactions, diet specializations, and others should be taken into account when interpreting current distribution patterns of *Limia* species.

## Introduction

Species distributions in natural systems are strongly modulated by climate, which ultimately affects both the ecology and physiology of organisms. This is particularly evident in ectothermic animals. Janzen (1967) published one of the most prominent papers in ecology that connected climatic variation across latitude and elevation, physiological adaptation and species distribution in a synthetic theory commonly referred as “Janzen’s hypothesis” (Ghalambor *et al*., 2006; Muñoz and Bodensteiner, 2019). One of the main predictions of this hypothesis is that due to the decrease in mean annual temperature with elevation, the seasonal temperature overlap is lower in the tropics than in temperate regions. Hence, mountain passes in the tropics may represent more effective physiological barriers to dispersal than the topographical component of change in altitude (Ghalambor *et al*., 2006). Therefore, the low overlap in temperature regimes between low and high elevations in the tropics should select for organisms with relatively narrow thermal tolerances. Janzen’s hypothesis also predicts that species develop physiological adaptations mirroring the range of ecological variation present in their surrounding area with populations living in high altitude evolving narrow tolerance for colder temperatures while low altitude populations developing narrow tolerance for warmer temperatures. Janzen’s hypothesis has been widely adopted and some studies have provided at least partial evidence at both local and global scale supporting his predictions and assumptions in both terrestrial and aquatic ectothermic organisms. For example, Pintanel *et al*. (2019) found that frog species occurring in open habitats, such as in valleys and lowland environments in general, had higher tolerance to high temperatures (CTmax) than species restricted to forest habitats, showing small climatic overlap across an elevation gradient. Moreover, Polato *et al*. (2018) provided strong evidence in support of Janzen’s hypothesis showing that tropical stream insects had noticeably narrower thermal tolerances and a lower dispersal ability than temperate species, which result in higher tropical speciation rates.

However, despite of general support of the theory, several components have never been thoroughly tested and critically evaluated across multiple taxa, potentially questioning the generality of Janzen’s theory (Ghalambor *et al*., 2006). In addition, under Janzen’s hypothesis is unclear whether the predictions refer to individual thermal niches or species thermal niches, which in fact are determined by different factors (Hua, 2016). In fact, some studies have not found support for Janzen’s theory: in amphibians (Valdivieso and Tamsitt, 1974) and in *Anolis* lizards from Hispaniola (Muñoz and Bodensteiner, 2019) factors such as daily variation in temperature and behavioral mechanisms might cause deviations from Janzen’s predictions. Furthermore, Navas *et al*. (2013) demonstrated that the effect of different microclimates within a specific biome is more relevant for species distributions than just the elevation at which certain species of amphibians may be found. McCain (2009) provided additional evidence for the effects of thermoregulation, daily temperature variability, and other climate variables such as precipitation as potential variables that could explain distribution ranges across multiple groups of vertebrates; including mammals, birds, reptiles and amphibians. In other words, Janzen’s theory may have to be amended by including more complexity.

Several key features related to the geographic distribution of *Limia* make these fishes an excellent system to explore how temperature fluctuations associated to elevational gradients might be linked to dispersal. *Limia* fishes are one of the most dominant groups in freshwater ecosystems in the Caribbean with at least 19 endemic species on Hispaniola and one endemic species each occurring in Cuba, Jamaica, and Grand Cayman (Burgess and Franz, 1989; Rodriguez, 1997; Hamilton, 2001; Rodriguez-Silva *et al*., 2020). These freshwater fishes occur in a wide distribution range occupying diverse aquatic habitats on these islands (Weaver *et al*., 2016 a). Although the altitudinal distribution of freshwater fish species in general is known to be considerably more constrained than in terrestrial species by several factors including for example productivity, physicochemical characteristics of the water and others (Jaramillo-Villa *et al*., 2010; Graham *et al*., 2014; Carvajal-Quintero *et al*., 2015), differences in altitudinal distribution in species of the genus *Limia* can be observed in natural habitats. In the present study, we tested some predictions of the Janzen’s hypothesis at the local scale through the analysis of the individual thermal niche breadth in several populations of livebearing fishes of the genus *Limia* and its relationship with their altitudinal distribution in some islands of the Greater Antilles in the Caribbean.

According to theory, we hypothesize that populations of species distributed in lowland habitats have evolved to resist higher extreme temperatures, which may be a factor limiting their dispersal into higher elevations. Conversely, populations occurring at higher elevations in mountain streams should have evolved to cope with lower temperatures, which reduce dispersal abilities into warmer habitats. Specifically, we predict that low elevation populations will be more tolerant to higher temperatures than mid and high elevation populations showing higher critical thermal maximum (CTmax) and critical thermal minimum (CTmin). In contrast, high elevation populations are expected to be more tolerant to lower temperatures showing lower CTmax and CTmin values. We also predict the thermal breadth (the range of temperatures they can tolerate) to be smaller for higher altitude fishes as result of little variability in CTmax.

## Materials and methods

The care and use of experimental animals complied with the University of Oklahoma animal welfare laws, guidelines and policies as approved by Animal Welfare Assurance on file with the Office of Laboratory Animal Welfare under the assurance number A3240-01. Experiments were performed under the approved IACUC protocol R17-011 and specimens were collected in the field as part of surveys of the native livebearing fishes of the Greater Antilles (protocol R18-005). No fishes were euthanized nor surgical procedures were performed.

### Study area and species

Even though thermal regimes of Greater Antillean streams are relatively stable, geological differences among islands lead to some climate heterogeneity that can generate environmental barriers. Hispaniola, for example, has several mountain ranges of more than 2000 meters in altitude. For instance, Pico Duarte in the Dominican Republic reaches 3098 meters and is the highest peak in the Caribbean. Previous studies have shown that high elevation specialists (mainly amphibians and reptiles) have evolved on Hispaniola as a consequence of this climate heterogeneity (Wollenberg *et al*., 2013; Muñoz *et al*., 2014). Mountains reaching around 2000 meters can be also found in eastern Cuba (Pico Turquino) and Jamaica (Blue Mountains) with significant levels of biodiversity associated.

In this study we analyzed populations of some species such as *L. perugiae, L. vittata, L. yaguajali* and *L. sulphurophila* that were reported to live in low elevation, warm environments including saline coastal lagoons. We also included in the analysis other species such as *L. zonata* and *L. melanogaster* that were obtained from low to intermediate elevations and often associated to relatively cool springs. Finally, one population of *L. dominicensis* and another of *L. versicolor* collected in mountain streams at relatively high elevations (see Table 1) were analyzed too. Specifically, we compared thermal breadth for the eight populations of the *Limia* species abovementioned that naturally occur in Cuba, Hispaniola, and Jamaica (Figure 1). For a more fine-grained picture, we analyzed three additional populations of *L. perugiae*, the most widely distributed species of *Limia* on Hispaniola in order to test for local adaptation to environmental variation in temperature.

**Figure 1:**
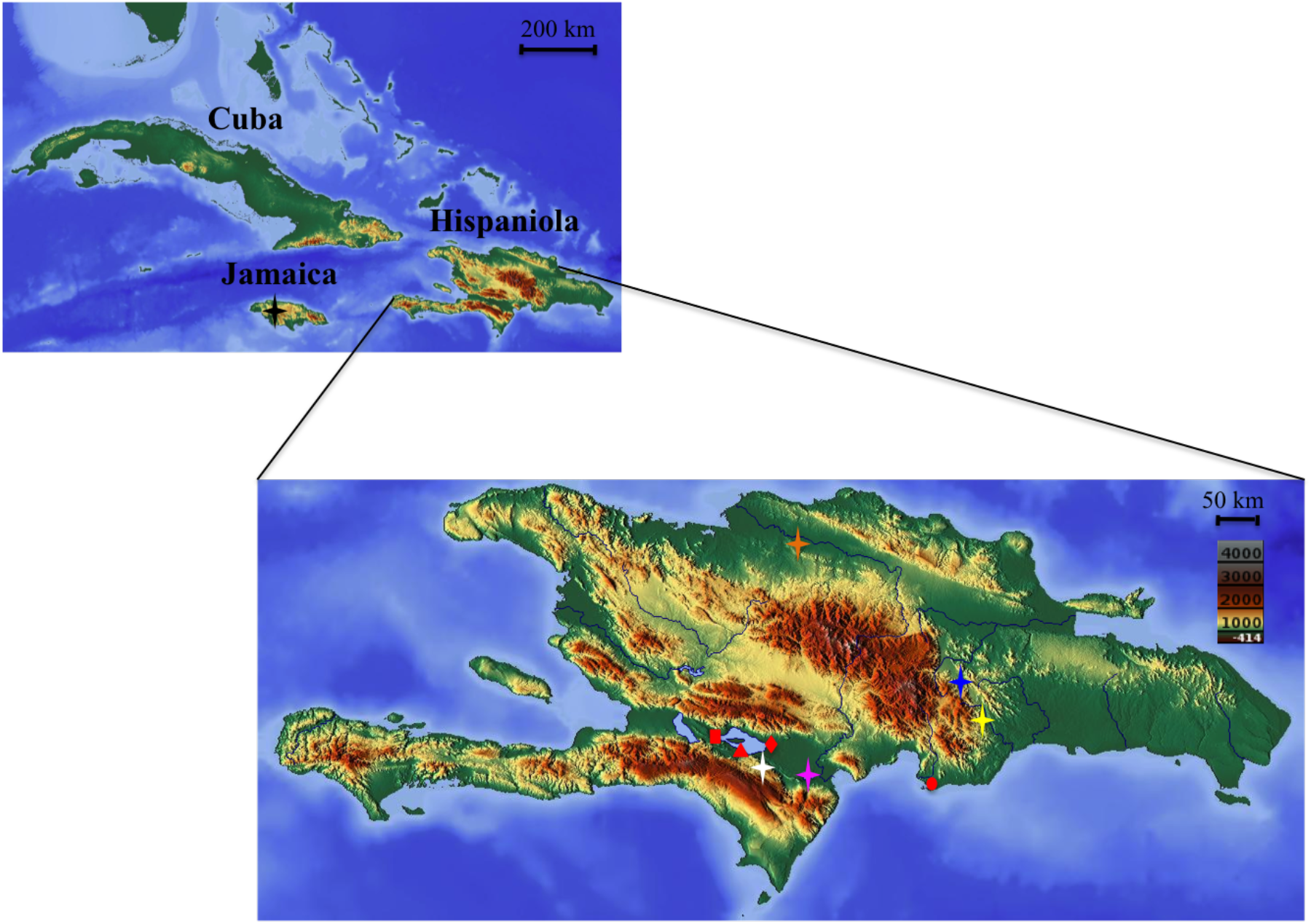
Geographic distribution of the populations of *Limia* (with known origins) analyzed in this study. Red rectangle: *L. perugiae* (Azufrada), Red triangule: *L. perugiae* (Lake Enriquillo), Red diamond: *L. perugiae* (La Zurza), Red circle: *L. perugiae* (Las Salinas), Orange star: *L. yaguajali*, Blue star: *L. zonata*, Black star: *L. melanogaster*, White star: *L. dominicensis*, Yellow star: *L. versicolor*, Pink star: *L. sulphurophila*.

**Table 1:** List of species and populations of *Limia* included in this study with GPS coordinates and elevation in meters above mean sea level (m.a.m.s.l.) of original collecting sites. Number of individuals analyzed of each sex is also included in all cases.

| **Species** | **Locality/GPS**<br>**coordinates** | **Collection**<br>**year** | **Sample size and**<br>**sex** | **Elevation**<br>**(m.a.m.s.l.)** |
| --- | --- | --- | --- | --- |
| *Limia perugiae* | Lake Enriquillo<br>18° 24' 4.61" N<br>71° 34' 16.61" W | 2014 | 7 females<br>3 males | 0 |
| *Limia perugiae* | Las Salinas (Bani)<br>18° 12' 43.488" N<br>70° 32' 27.095" W | 2011 | 6 females<br>4 males | 0 |
| *Limia perugiae* | Azufrada<br>18° 33' 40.212" N<br>71° 41' 50.928" W | 2003 | 5 females<br>5 males | 0 |
| *Limia perugiae* | La Zurza<br>18° 23' 56.08" N<br>71° 34' 13.22" W | 2014 | 5 females<br>5 males | 0 |
| *Limia vittata* | Cuba<br>(Unknown coordinates) | Unknown | 5 females<br>5 males | Unknown |
| *Limia sulphurophila* | Cabral (Barahona<br>Province)<br>18° 14' 45.902" N | 2014 | 6 females<br>4 males | 26 |
|  | 71° 13' 23.944" W |  |  |  |
| *Limia yaguajali* | Arroyo del Agua (Jamao<br>al Norte)<br>19° 37' 49.872" N<br>70° 26' 58.056" W | 2018 | 5 females<br>5 males | 46 |
| *Limia melanogaster* | Roaring River<br>18° 17' 00.0" N<br>78° 03' 22.0" W | 2017 | 9 females<br>3 males | 71 |
| *Limia zonata* | Rio Yuna (Bonao)<br>18° 57' 33.1" N<br>70° 24' 32.06" W | 2014 | 6 females<br>4 males | 158 |
| *Limia versicolor* | Rio Basima<br>18° 42' 5.605" N<br>70° 11' 50.387" W | 2018 | 5 females<br>5 males | 181 |
| *Limia dominicensis* | Puerto Escondido<br>18° 19' 6.93" N<br>71° 34' 14.24" W | 2014 | 5 females<br>5 males | 399 |

All populations analyzed but *L. vittata* came from wild caught stocks that were transported into the United States and then kept in common garden conditions at the University of Oklahoma for variable periods of time (Table 1). *L. vittata* specimens were obtained from aquarium stocks and have been kept in common garden conditions at a greenhouse in the Aquatic Research Facility at the University of Oklahoma for more than 10 years.

### Laboratory methods

We used the critical thermal method (Cowles and Bogert, 1944) to describe variation in temperature tolerance in 113 adult fish of eight *Limia* species representing a total of 11 different populations (Figure 1). Prior to testing all fishes were acclimated to laboratory conditions with temperatures ranging between 25^0^C-27 ^0^C for 45 days. This was the most common temperature range for the species at their origins. We tested 10 reproductively mature adult fish of both sexes per population.

Fishes were individually tested under a constant increase (to determine critical thermal maximum, CTmax) or decrease (to determine critical thermal minimum, CTmin) in temperature until reaching an appropriate endpoint. The endpoint we used to determine CTmax was the pre-death thermal point at which signals of sudden onset of muscular spasms appeared (Lutterschmidt and Hutchison, 1997; Beitinger *et al*., 2000). In the case of the CTmin the endpoint used was the absence of motion of the pectoral fins in which the fish did not start to move again even when the experimenter disturbed the fish (Fischer and Schlupp, 2009). Using this data, temperature tolerance was calculated as the arithmetic means of high and low temperatures (CTmax or CTmin) at which the endpoint was reached by individuals in the sample (Lowe and Vance, 1955). We also calculated the thermal breadth as CTmax minus CTmin for each individual. The three measures, although connected, reflect different physiological properties of the species.

Before the actual experiment fishes were not fed for 24 hours. Each fish was individually tested in a spherical 2-liter glass container. After a 10-minutes acclimation period before each trial, the container was heated using a concave heating plate at a constant rate of 1^0^C/min while temperature was constantly monitored with a thermometer in order to record CTmax. Each trial was immediately stopped, and the final temperature was measured once the fish showed symptoms of sudden muscular spasms, which were characterized by disorganized and high frequency muscular movements. All fish were weighed after each trial, placed in individual tanks and allowed to rest for at least 72 hours until application of the other extreme temperature to the same individual.

Temperature exposure (CTmax or CTmin) that a fish experienced first was randomized to avoid an order effect. This was also evaluated statistically.

A similar procedure was followed to test the fish’s tolerance to cold temperatures or CTmin. In this case, we placed the fish in a similar 2-liter glass container and after a 10-minute acclimation period, we continuously added cold water of 3^0^C −4^0^C to the system for a rate of temperature change of 1^0^C/min. CTmin was measured at the point in which fish showed total absence of movement. No mortality was associated with the trials and after the experiments all fishes were returned to their respective stock tanks

### Climate index overlap

Monthly maximum and minimum temperatures for the 2010-2019 period were extracted from WorldClim database for the following localities included in table 1: La Zurza, Cabral, Arroyo del Agua, Roaring River, Rio Yuna, Rio Basima and Puerto Escondido. We calculated the average monthly temperature for each site during the last 10 years in order to measure pairwise thermal overlap between each focal site and the other localities using Janzen’s (1967) equation:

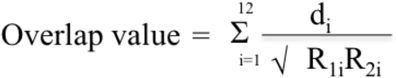

where d_i_ is the thermal overlap between the focal site and each other site for the i^th^ month or the amount (in Celsius degrees) of one thermal regime that is included within the other, R_1i_ is the difference between the monthly mean maximum and minimum for the focal site and R_2i_ is the corresponding value for each other site of the study. As temperature overlap increases the overlap value increases up to a value of 12 which is the point where thermal regimes between two sites share identical monthly maximum and minimum temperatures throughout the year.

### Phylogenetic signal

A common caveat of studies like this is that any pattern found might not necessarily reflect adaptations but be due to species relatedness. To test for this we used a recent phylogeny (Weaver *et al*., 2016 b) and conducted a test of phylogenetic signal to assess whether correlations in temperature tolerance among species may be due to their shared evolutionary history or to other factors (Gingras *et al*., 2013; Kamilar and Cooper, 2013; Gilbert *et al*., 2018; Arnaudo *et al*., 2019). For this analysis we used Pagel’s lambda (λ) (Pagel, 1994) as a quantitative measure of this relationship. The Pagel’s λ has been shown to be a very robust indicator of a correlation between ecological and evolutionary processes even for incompletely resolved phylogenies (Molina-Venegas and Rodriguez, 2017; Leiva *et al*., 2019). We based this analysis on the phylogeny published by Weaver *et al*. (2016 b), which includes most of the species used in this study.

### Data analysis

For data analysis of thermal tolerance, we used one-way ANOVA’s, after confirmation of homogeneity of variances by Levene’s tests and normality by Shapiro and Wilk’s tests. We used three different ANOVA’s to compare the mean CTmax, CTmin and temperature ranges (breadth) among eight different populations of *Limia*. Scheffe’s post hoc tests were used to make comparisons between groups to distinguish populations that differed from others in extreme thermal tolerance or temperature ranges. The lack of order effect was statistically confirmed (p>0.05) using t-test analyses to compare CTmax and CTmin means of fish that were tested CTmax then CTmin versus fish that were tested CTmin then CTmax.

All statistical analyses were performed in SPSS 23. We performed independent ANOVA analyses because we considered CTmax and CTmin as ecologically and evolutionary independent variables with potentially different adaptive benefits. Similar approaches that consider the effects of these variables (and also acclimation temperature) as independent have been used in other studies examining thermal tolerances in ectothermic animals (Spotila, 1972; Layne and Claussen, 1982). To further explore a potential role for local adaptation in thermal tolerance within a widely distributed species, we also compared the same variables through separate ANOVA analyses in four populations of *L. perugiae*. In order to calculate climate index overlap we used the R package raster (Hijmans, 2020) to extract minimum and maximum monthly temperatures values to our sampled locations. Finally, analysis of phylogenetic signal was computed using the R package (R Core Team, 2013) phytools (Revell, 2012).

## Results

### Analysis of phylogenetic signal

We performed analyses of phylogenetic signal using a previously-inferred phylogeny based on three mitochondrial (12S, ND2, Cytb) and two nuclear (MYH6, Rh) genes (Weaver *et al*., 2016 b) to determine whether more similar values in CTmin, CTmax and thermal range were associated with more closely related species more often than expected by chance. None of the analyses found a significant effect of phylogeny in explaining thermal breadth among the *Limia* species studied: CTmin (λ=6.257973e-05, p=1.000), CTmax (λ =0.4056982, p=0.78255460) and thermal range (λ=1.16242, p=0.362986) (Supplementary Information Figure 1).

### Analysis of climate index overlap

Overall, thermal overlap along altitudinal gradients among the study locations was relatively high but there was still some variation across pair of sites (Table 2). Climate index overlap decreased as the differences in elevations were more conspicuous. Particularly, Puerto Escondido (the highest collecting site included in the analysis) showed the lowest climate overlap with all other sites, which indicates that there are some differences in habitat temperatures.

**Table 2:** Pairwise values of climate index overlap across the seven collecting sites showing different altitudinal gradients. Climate index overlap can take values up to a maximum of 12 where the higher values indicate more temperature overlap.

|  | La Zurza | Cabral (Barahona) | Jamao al Norte | Roaring River | Rio Yuna (Bonao) | Rio Basima | Puerto Escondido |
| --- | --- | --- | --- | --- | --- | --- | --- |
| La Zurza | - |  |  |  |  |  |  |
| Cabral (Barahona) | 11.47 | - |  |  |  |  |  |
| Jamao al Norte | 10.63 | 11.15 | - |  |  |  |  |
| Roaring River | 10.81 | 11.22 | 10.28 | - |  |  |  |
| Rio Yuna (Bonao) | 10.29 | 11.39 | 11.59 | 9.96 | - |  |  |
| Rio Basima | 10.05 | 10.55 | 11.74 | 9.65 | 11.84 | - |  |
| Puerto Escondido | 8.93 | 9.38 | 10.17 | 8.45 | 10.57 | 10.79 | - |

### Inter-specific analysis of thermal tolerance

The overall range of temperature tolerance for the eight species included in the analysis was 12^0^C (CTmin) to 41.2^0^C (CTmax), which may be considered as a broad range for tropical fishes when considering the overall climatic stability present in the tropics in terms of temperature fluctuations. A one-way ANOVA was conducted to compare the effect of varying distribution according to elevation on the temperature tolerance under CTmin and CTmax conditions. There were significant differences in thermal limits for both CTmin (One-way ANOVA, F (7, 72) = 41.977, p<0.001) and CTmax (One-way ANOVA, F (7, 72) = 14.878, p<0.001) among species after testing 80 individuals. The highest temperature tolerance was recorded for *L. sulphurophila* with an average CTmax of 40.9^0^C. This species also showed the lowest tolerance to low temperatures with an average CTmin of 16.7^0^C, which might suggest this species could be adapted to live in warmer habitats. A post hoc analysis also showed that *L. sulphurophila* differed significantly from all other species in both CTmin and CTmax (Scheffe, p<0.05) except for *L. zonata* in CTmin (Scheffe, p=0.137) (Figure 2).

**Figure 2:**
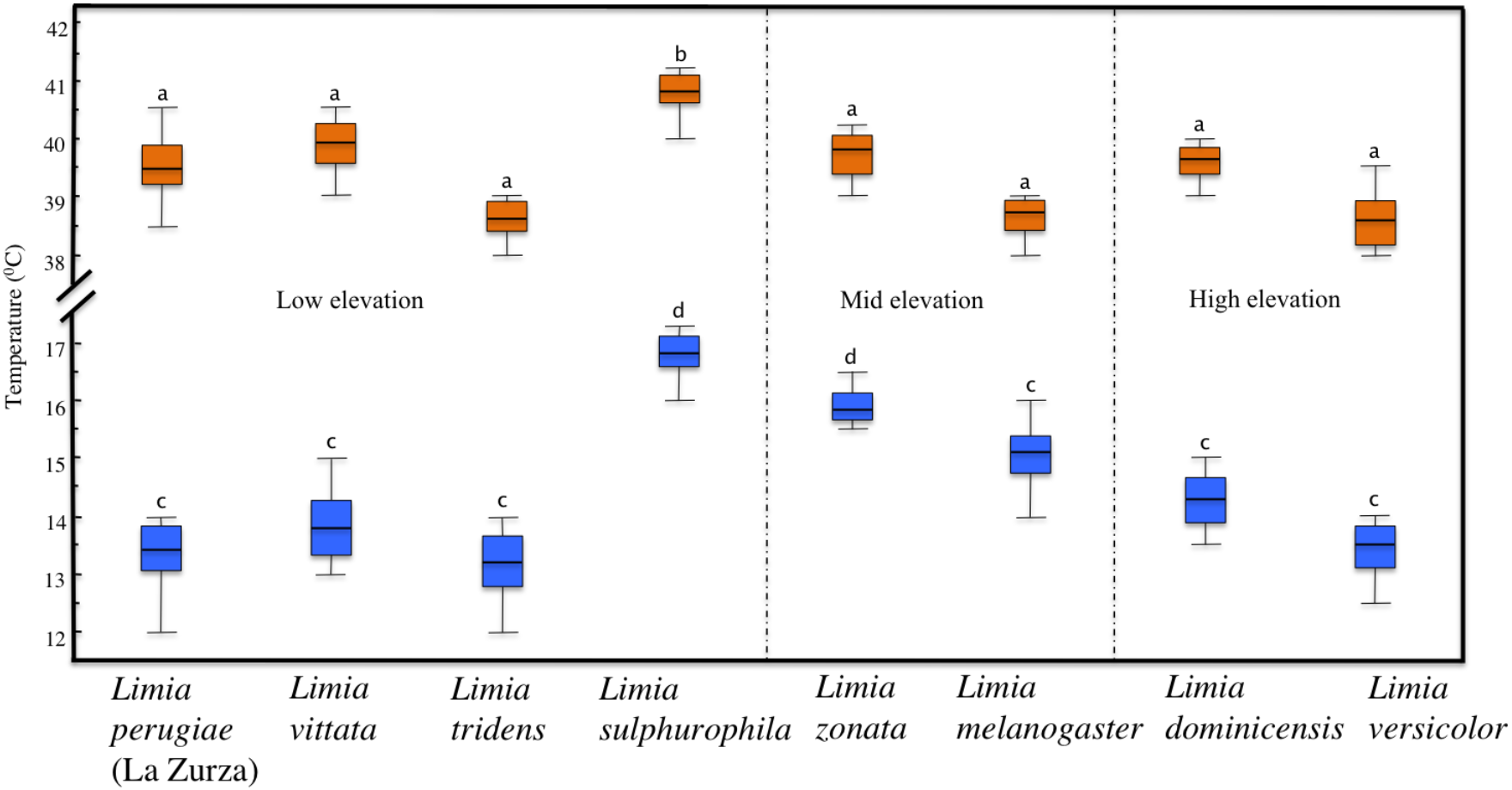
Thermal tolerances of the eight species of *Limia* included in this study. Species are grouped according to their elevation distribution in three groups: low elevation species (left), mid elevation species (center) and high elevation species (right). Each box plot represents the median, interquartile ranges, maximum and minimum values of either CTmin or CTmax for each species. CTmin is shown by blue box plots and CTmax by orange box plots.

Another ANOVA was used to compare temperature ranges among populations. The analysis showed significant differences (One-way ANOVA, F (7, 72) = 15.993, p<0.001). *L. melanogaster* displayed the narrowest range of thermal tolerance, which differed from all other populations (Scheffe, p<0.05) but not from *L. sulphurophila* (Scheffe, p=1.000), *L. zonata* (Scheffe, p=0.994) and *L. versicolor* (Scheffe, p=0.117). Our data suggested that the most tolerant species to extreme temperatures were *L. perugiae*, *L. yaguajali*, and *L. vittata* (species distributed in low elevations) followed by *L. versicolor* and *L. dominicensis* (species which distribution extends into much higher elevations) since these two groups of species displayed broader ranges of temperature tolerance (Figure 3) and their ranges did not differ significantly from each other (Scheffe, p>0.05).

**Figure 3:**
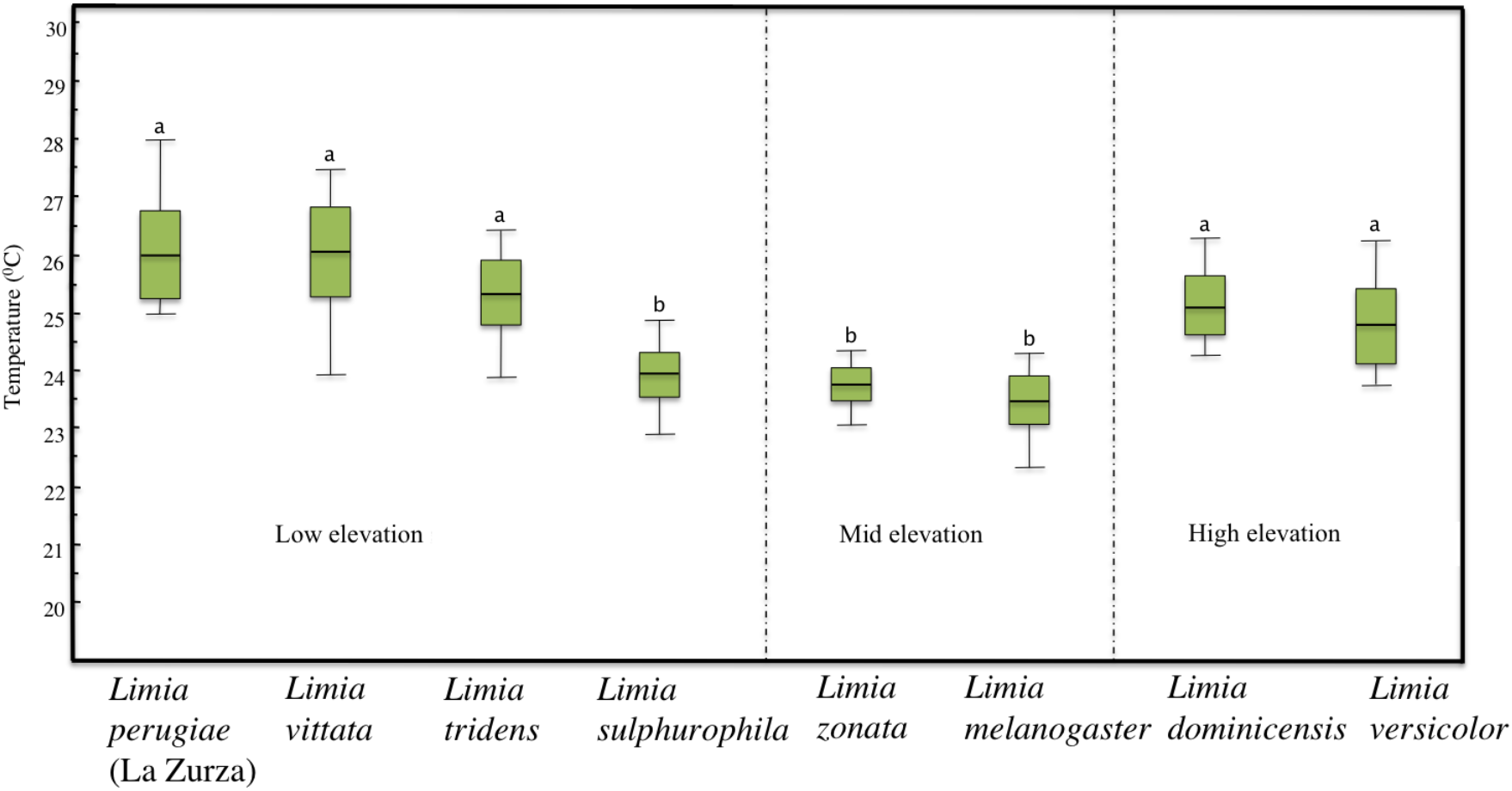
Box plots of the thermal ranges of the eight species of *Limia* included in this study. Species are grouped according to their elevation distribution in three groups: low elevation species (left), mid elevation species (center) and high elevation species (right). Each box plot represents the median, interquartile ranges, maximum and minimum values of either CTmin or CTmax for each species.

### Intra-specific analysis

Population analyses for *L. perugiae* showed significant differences in CTmax (One-way ANOVA, F (3, 36) = 6.118, p=0.02) and CTmin (One-way ANOVA, F (3, 36)= 20.982, p<0.001). The population from Lake Enriquillo differed from the other three in both CTmax (Scheffe, p<0.05) and CTmin (Scheffe, p<0.001) (Figure 4). There were also significant differences in ranges of thermal tolerance among populations of *L.perugiae* (One-way ANOVA, F (3, 36) = 3.409, p=0.028). In this case the population from Lake Enriquillo had significant differences with the population from Azufrada (Scheffe, p=0.037). This is surprising as the Azufrada population is also from Lake Enriquillo, just the north shore, not the south shore.

**Figure 4:**
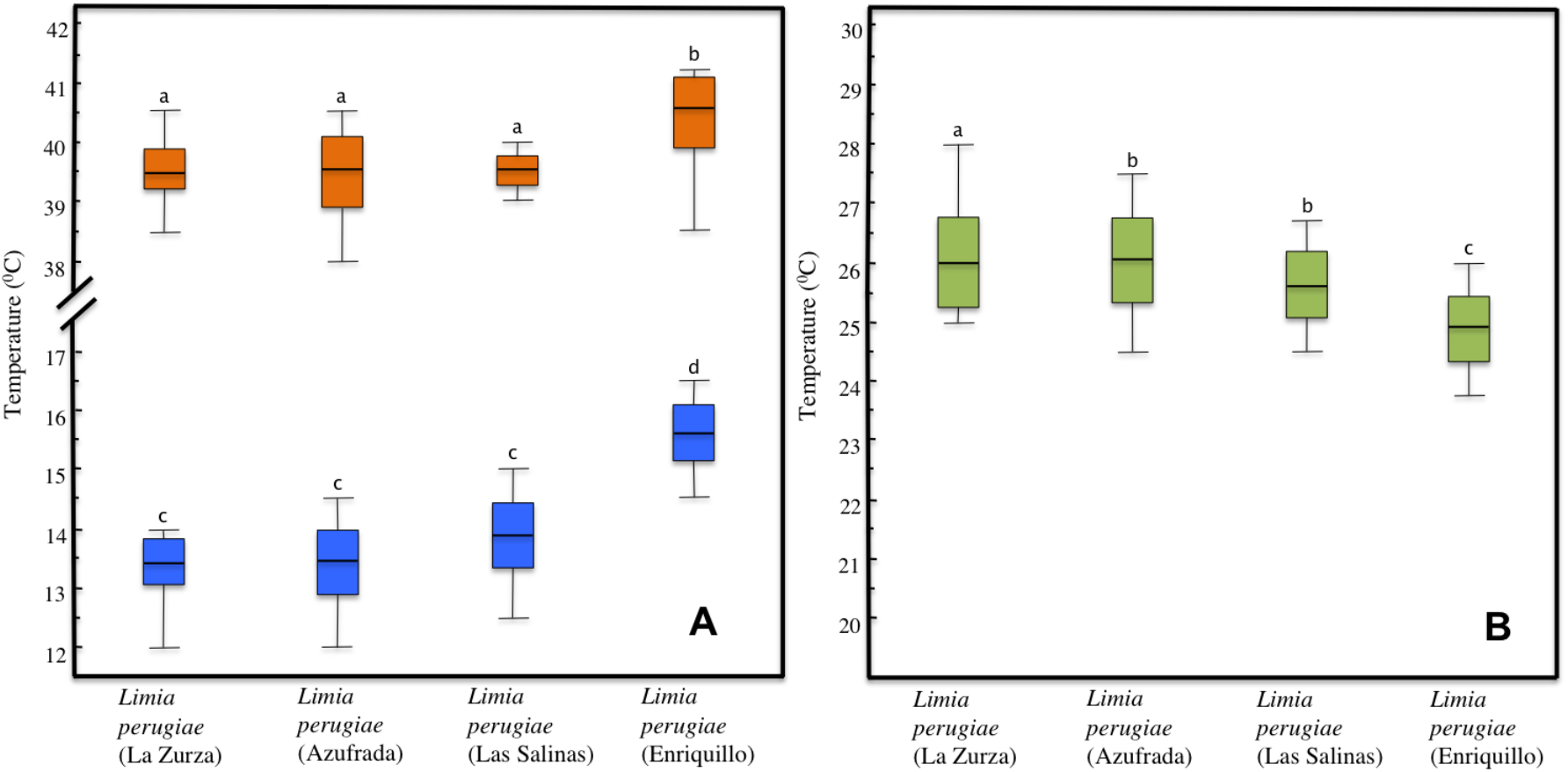
Thermal tolerances of four populations of *L. perugiae* showing CTmin in blue box plots and CTmax in orange box plot (A), and box plots of thermal ranges of the four populations analyzed (B). Each box plot represents the median, interquartile ranges, maximum and minimum values.

## Discussion

Our evidence suggests that thermal tolerance and altitudinal distribution of the populations of *Limia* species analyzed in this study are not be related to temperature gradients expected in nature. The species studied here showed thermal tolerances not predicted by Janzen’s hypothesis. Generally, even species from high altitudes (for the tropics) have broad thermal tolerances similar to species distributed in low elevations. Hence, the observed pattern does not separate the species as predicted.

Additionally, phylogeny did not explain species relationships according to thermal tolerance. However, failure to detect statistical significance may be due to the rather low number of species included in the analyses. Small sample sizes have been shown to influence the uncertainty and the expected values of most indices of phylogentic signal, including the Pagel’s lambda (λ) (Munkemueller *et al*., 2012). Powerful phylogenetic comparative analyses typically demand trait and phylogenetic data for over 50 species and *Limia* only has 22 described species, which simply cannot satisfy the sampling requirements for robust macroevolutionary inferences.

Temperature tolerance ranges have been shown to shape species distributions and community compositions for some ectotherms in both tropical and temperate climates. Snyder and Weathers (1975) offered experimental evidence on the close relationship between the range of temperature tolerance and the environmental temperature variation in the distribution of several species of amphibians, showing that an increase in the environmental temperature variation also increases the range of temperature tolerance and consequently the distribution range of species. Estimating temperature tolerances and temperature ranges through the analysis of lower (CTmin) and upper (CTmax) thermal limits has been shown to be an efficient and useful method to assess species’ capacity to acclimate to temperature changes in several ectotherms including terrestrial species (i.e. arthropods, reptiles and amphibians) and aquatic organisms (i.e. arthropods, mollusks and fish) (Van Berkum, 1988; Sunday *et al*., 2011; Buckley and Huey, 2016). However, when using this methodology, the results can be influenced by experimental protocols and conditions in which either CTmin or CTmax are measured. Factors such as acclimation temperature of individuals being tested, the cooling or heating rate used, and also the non-lethal endpoint chosen by the experimenter to determine CTmin or CTmax can influence the results (Lutterschmidt and Hutchison, 1997). Fortunately, though, the method has been in use for a long time, which has allowed testing and standardizing protocols for different animal groups. Hence, the technique offers repeatable and rapid quantitative measure of the thermal limits as well as predicts optimal temperate ranges of multiple species (Fischer and Schlupp, 2009; Kingsolver and Umbanhowar, 2018). Although specimens used in this experiment were kept in common garden conditions for different lengths of time, which might influence their tolerance to critical thermal limits; we standardized the acclimation time to a specific temperature range (25^0^C −27^0^C) for 45 days under laboratory conditions. This acclimation time is considerably longer than others previously reported in studies of thermal limits in ectothermic organisms (Chanthy *et al*., 2012; Moyano *et al*., 2017; Tongnunui and Beamish, 2017), which ensures our data truly reflects the actual thermal tolerances of the species.

Janzen (1967) predicted that species occurring in high altitude in the tropics are specialized for lower temperatures than are low altitude species, which should be better adapted to cope with warmer temperatures. However, our results do not offer support for this prediction suggesting that factors other than temperature shape the distribution in populations of livebearing fishes of the genus *Limia*. In general, species and populations widespread in lowland habitats (*L. perugiae* populations from Hispaniola and *L. vittata* from Cuba) seem to be very tolerant to extreme temperatures (CTmax and CTmin) suggesting that they could also live in mountainous habitats. Given this, what could explain that none of these two species are found in high elevation streams either in Cuba or Hispaniola? We suggest that this is probably due to biotic factors such as competition. In the case of *L. vittata* in Cuba other dominant livebearing fishes (genera *Girardinus* and *Gambusia*) exploit available niches in mountain streams and on Hispaniola other *Limia* species and also species of *Poecilia (P. dominicensis, P. hispaniolana* and *P. elegans*) seem to be restricting *L. perugiae* to lowland environments. Evolutionary trade-offs between broad tolerance and competitive habitats have been shown to be common in different taxa. For instance, Robinson and Terborgh (1995) showed that interspecific aggression more that habitat suitability might explain spatial segregation patterns observed in Amazonian birds, and Griffis and Jaeger (1998) defined interspecific competition as cause of extinction of a species of salamander (*Plethodon shenandoah*) in the mountains of Shenandoah National Park, Virginia, USA.

Conversely, among species with distribution ranges that mostly include mid-elevations, there is a less consistent pattern in tolerance to extreme temperatures but with a general trend towards lower ranges of tolerance. Such are the cases of *L. melanogaster* from Jamaica and *L. zonata* from Hispaniola, which showed low tolerance ranges and were particularly sensitive to low temperatures. Our recent exploratory work in the Caribbean has recorded these two species associated with permanent freshwater springs and spring runs that buffer temperature fluctuations. The two species abovementioned seem to be physiologically adapted to relatively narrow fluctuations in temperature, which may explain their limited tolerance range.

Another result of our study that runs counter to Janzen’s hypothesis is that two strictly mountainous and locally distributed populations of the species *L. versicolor* and *L. dominicensis*, exhibited relatively broad tolerance ranges similar to species that typically are widespread distributed in low elevation environments. Increasing altitude is always accompanied by a decrease in annual average temperature (Sarmiento, 1986), which may indicate high elevation organisms are better adapted to cope with low temperatures. This general pattern was also present in our analysis where the highest study site, in this case Puerto Escondido with one population of *Limia dominicensis*, exhibited the lowest climate overlap index with the rest of the other sites. However, in tropical ecosystems species living at high elevations might benefit from evolving broad thermal tolerance to deal with diurnal changes in temperature (Ghalambor *et al*., 2006), which in turn might explain why the two species also show a moderate tolerance for high temperatures. Temperature tolerances of high and mid elevation populations of *Limia* species relate to results of other studies in tropical amphibians (Navas, 1996; Navas *et al*., 2013), which showed that species occurring in intermediate elevations were likely stenothermic given the relative thermal stability of those habitats. Conversely, species from higher elevations seemed to have evolved to lead with more changeable temperature (differences between day and night temperatures) and consequently develop broader tolerance ranges.

Our results also provide insights into local physiological adaptations of thermal tolerance within species, which suggests that conspecific populations in diverse habitats have somewhat independent evolutionary pathways (Snyder and Weathers, 1975). First, *L. perugiae* from the south shore of Lake Enriquillo, near where *L. sulphurophila* is found differed from other three populations by showing a narrower thermal breadth. This result – together with the lack of phylogenetic signal in thermal tolerance for species – emphasizes the importance of biogeographical processes more than just phylogenetic patterns in analyses of climatic niche (Coelho *et al*., 2019). Second, *L. sulphurophila*, a locally distributed species mainly known from sulfur springs on the southeastern shore of Lake Enriquillo in the Dominican Republic (Rivas, 1980), seems to be locally adapted to live in a high temperature sulfidic environment and is also physiologically adapted to resist high temperatures. This may prevent competitive exclusion as shown in fish species from temperate climates (Ohlberger *et al*., 2008).

Our study has implications for conservation and is also pertinent in the context of climate change and species resilience to short-term temperature spikes. Our data provide evidence of species and populations that would be more vulnerable to temperature variation. In this case, the ones occurring in cool permanent freshwater springs in mid elevations and also *L. sulphurophila* (a local endemic species) seemed to be more susceptible to temperature fluctuations because of their narrower thermal breadth. Another issue that may affect conservation of native *Limia* species is the introduction of invasive livebearing fishes, such as *Poecilia reticulata* (Guppy). This species has recently been reported as one of the most tolerant ornamental fish to extreme temperatures (Yanar *et al*., 2019), which may be additional evidence of the invasive success of guppies and in some extend explain why this species becoming dominant in tropical ecosystems.

The implications of temperature for fish physiology and fitness (Niehaus *et al*., 2012; Payne *et al*., 2016) make the analysis of thermal limits particularly important in determining distribution of fishes (Culumber *et al*., 2012). Even though our study does not provide a comprehensive test for Janzen’s hypothesis, it presents evidence at local scales to analyze how elevational gradients may affect the distribution of freshwater fishes, which is a barely studied zoological group in the Caribbean. While it does not offer evidence supporting Janzen’s predictions about climatic variation across elevations, physiological adaptation and species distribution for this group of fish; the study emphasizes the importance of testing the validity of Janzen’s mountain passes hypothesis across multiple taxa. In addition, like previous studies this work stresses the significance of other factors such as species interactions, diet specializations, and even thermoregulatory behavior (as shown by Muñoz and Bodensteiner (2019) in Caribbean anoles) when interpreting current altitudinal distribution patterns of species.

## Acknowledgements

We would like to thank the governments and corresponding ministries of Jamaica and the Dominican Republic for kindly issuing colleting permits. In addition, we thank to the Museo Nacional de Historia Natural “Prof. Eugenio de Jesus Marcano” in Santo Domingo, Dominican Republic and especially to Patricia Torres Pineda and Carlos Suriel. We also thank Carlos Rodriguez from the Ministry of Education in the Dominican Republic for his support. Thanks to Ricardo Betancur and Emanuell Ribero their insights and for helping with the analysis of phylogenetic signal. Thanks to Caryn Vaughn, Laura Stein and Bruce Hoagland for their advice and comments on the manuscript. We are grateful to Carolyn Burt, Kerri-Ann Bennett, Stephan Bräger, Heriberto Encarnación Lara, Kenia Ng Alvarado, and Marcos José Rodriguez for help with fieldwork. Trai Spikes, Sophie Huebler, Margaret Zwick, Zeeshawn Beg, and Nabiha Ahmad helped with fish care. Gabriel Costa helped with statistical analysis. Finally, we are grateful to the reviewers for constructive comments on the manuscript.

## Contributions

R. R. S. – Contributed to this manuscript with ideas, data generation (including fieldwork), data analysis, manuscript preparation and funding.

I. S. – Contributed to this manuscript with ideas, data collection in the field and manuscript preparation.

