## Supplemental Figure 1 for "Elevational gradients do not affect thermal tolerance at local scale in populations of livebearing fishes of the genus *Limia* (Teleostei, Poeciliidae)"

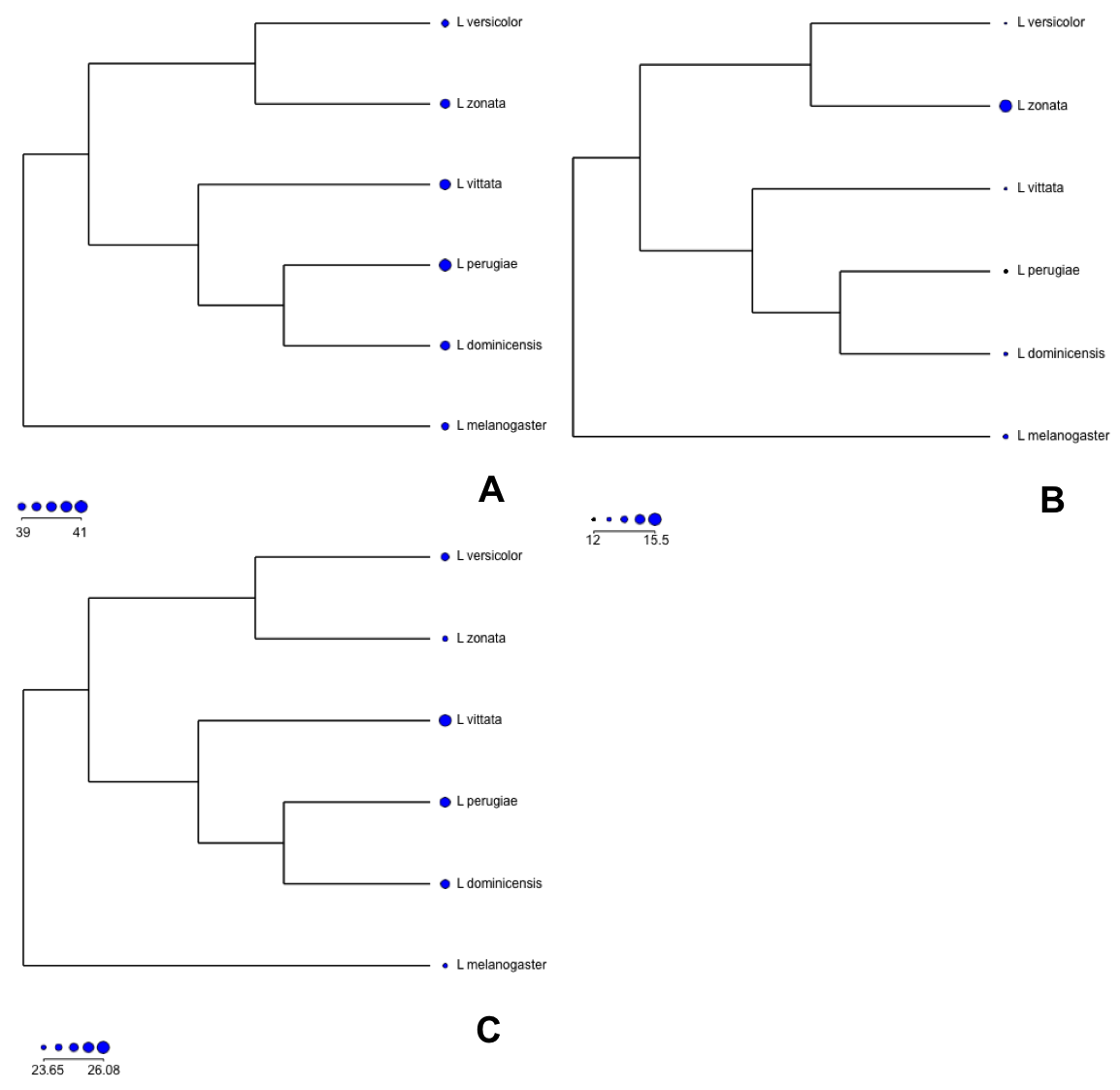

Supplementary Information Figure 1: Analysis of phylogenetic signal using Pagel’s lambda (λ). The phylogeny of species is based on mitochondrial (12S, ND2, Cytb) and nuclear (MYH6, Rh) gene sequences (Weaver *et al*., 2016 b). Circles at the tips of the phylogeny are sized in proportion to the average value of similarities in CTmax (A), CTmin (B) and thermal range (C) for each species. The legends in the figure also indicate the minimum and maximum values for each variable.
